# Size-reduced embryos reveal a gradient scaling based mechanism for zebrafish somite formation

**DOI:** 10.1101/211615

**Authors:** Kana Ishimatsu, Tom W. Hiscock, Zach M. Collins, Dini Wahyu Kartika Sari, Kenny Lischer, David L. Richmond, Yasumasa Bessho, Takaaki Matsui, Sean G. Megason

## Abstract

Little is known about how the sizes of animal tissues are controlled. A prominent example is somite size which varies widely both within an individual and across species. Despite intense study of the segmentation clock governing the timing of somite generation, how it relates to somite size is poorly understood. Here we examine somite scaling and find that somite size at specification scales with the length of the presomitic mesoderm (PSM) despite considerable variation in PSM length across developmental stages and in surgically size-reduced embryos. Measurement of clock period, axis elongation speed, and clock gene expression patterns demonstrate that existing models fail to explain scaling. We posit a “clock and scaled gradient” model, in which somite boundaries are set by a dynamically scaling signaling gradient across the PSM. Our model not only explains existing data, but also makes a unique prediction that we experimentally confirm—the formation of periodic “echoes” in somite size following perturbation of the size of one somite. Our findings demonstrate that gradient scaling plays a central role both in progression and size control of somitogenesis.

**Summary statement:** By comparing patterning in zebrafish embryos of different size we show that a dynamically scaling gradient in the presomitic mesoderm regulates somite size control.

## Main

Scaling― matching organ size to body size― is a fundamental property of developing organisms. Even within the same species, developing embryos often vary in size, due to environmental and maternal variability. In addition, embryo size can change drastically across developmental stages. Nevertheless, embryos robustly develop with invariant proportions, suggesting that some mechanism of pattern scaling is encoded in the developmental program (Cooke, 1981). While the scaling of morphogen-based patterning has received significant attention, both theoretically and experimentally (Ben-Zvi and Barkai, 2010; Gregor et al., 2005; Gregor et al., 2008; Inomata et al., 2013; Lander et al., 2011; McHale et al., 2006; O’Connor et al., 2006), understanding has been limited for scaling of other patterning processes, such as somite segmentation.

During embryogenesis, somites provide the first body segments in vertebrates, eventually giving rise to tissues such as the vertebrae and axial skeletal muscles. Somite segmentation occurs sequentially in an anterior to posterior progression along the presomitic mesoderm (PSM), with temporal and spatial periodicity. Temporal periodicity (e.g. somites are formed in symmetric pairs every 25 min in zebrafish (Schroter et al., 2008)) is known to be generated by a system of coupled cellular oscillators (Delaune et al., 2012; Lauschke et al., 2013; Masamizu et al., 2006; Palmeirim et al., 1997), called the segmentation clock, which is driven and synchronized by complex signaling networks (Dequeant et al., 2006; Hubaud and Pourquie, 2014; Krol et al., 2011). Yet, how these oscillations relate to the spatially periodic pattern of the mature somites and how somite sizes are determined remains controversial (Akiyama et al., 2014; Cooke and Zeeman, 1976; Cotterell et al., 2015; Lauschke et al., 2013; Shih et al., 2015; Soroldoni et al., 2014; Takahashi et al., 2010; Tsiairis and Aulehla, 2016).

Somites were first documented to scale in *Xenopus* following surgical bisection of the egg; the resulting embryos have smaller somites but the same number when compared to intact control embryos (Cooke, 1975). Although this experiment was performed more than 40 years ago, the underlying mechanism for somite scaling has not been identified. In particular, the relationship between PSM length and somite size has been disputed: previous groups have reported that in intact developing embryos, somite size does not scale with PSM size (Gomez et al., 2008), while in ex vivo culture of PSM, somite length has been shown to linearly scale with PSM length (Lauschke et al., 2013).

In this study, we demonstrate that somite length indeed scales with PSM length and that gradient scaling underlies somite scaling, using both surgically size-reduced and normally developing zebrafish embryos, in combination with live imaging, quantitative measurement, and mathematical modeling. We demonstrate that the inconsistency in the reported relationship between PSM size and somite size can be attributed to the time delay between somite specification and morphological boundary formation. We experimentally measured this delay and found that somite length always scales with PSM length when this delay is considered. This result led us to evaluate several variables that could potentially modulate somite length. We found that clock period, axis elongation speed, and clock gene expression patterns did not scale, whereas the Fgf activity gradient did scale with PSM length. Given this observation, we developed a “clock and scaled gradient model” based on the original clock and wavefront model (Cooke and Zeeman, 1976) with a simple yet important refinement; in our model, the gradient responsible for setting wavefront position dynamically scales to the size of the PSM. Using transplants, we show that somite derived signals can inhibit Fgf signaling providing a potential mechanism for gradient scaling. The clock and scaled gradient model not only explains existing experimental data but also inspired a novel experimental test with an unintuitive outcome—the creation of “echoes” in somite size following perturbation of the system. Together, we present the quantitative study of somite scaling as an experimental platform to test the feasibility of multiple theoretical models.

## Results

### Somite length at specification scales with PSM length throughout developmental time

Although somite length has been shown to scale with overall body length in *Xenopus* (Cooke, 1975), whether somite length scales with PSM size has been controversial (Gomez et al., 2008; Lauschke et al., 2013). To test this relationship we measured somite length and PSM length using live imaging. Initially we did not observe a clear relationship between PSM length and somite size (see Fig.1F, without delay). However, somite specification within the PSM occurs long before the appearance of the morphological boundaries (Akiyama et al., 2014; Bajard et al.,; Dubrulle et al., 2001; Elsdale et al., 1976; Giudicelli et al., 2007; Ozbudak and Lewis, 2008; Primmett et al., 1989; Roy et al., 1999) (Fig. 1A), and thus we speculated that the inconsistency with respect to somite scaling could be attributed to this delay. Although previous studies have shown the delay is around 4-5 cycles, the delay duration could vary along developmental stages. To examine if somite length scales with PSM length when this specification to formation delay is considered, we experimentally measured this delay using embryos from different developmental stages. Dual-specificity phosphatase inhibitor BCI is known to act immediately on Fgf signaling leading to an eventual reduction of somite size (Fig. S1) (Akiyama et al., 2014). We transiently treated embryos at 5 somite stage (ss), 10ss, and 15ss with BCI and measured the length of the newly formed somites using live imaging for six subsequent cycles (Fig. 1B and C). Regardless of the developmental stage for the pulse BCI treatment, we observed 4-cycle delay on average before a visibly smaller somite formed (Fig. 1D). Our experimentally determined delay is similar, albeit slightly shorter, to what has been proposed in previous work (4-5 cycles) (Akiyama et al., 2014; Bajard et al.,; Dubrulle et al., 2001; Elsdale et al., 1976; Giudicelli et al., 2007; Ozbudak and Lewis, 2008; Primmett et al., 1989; Roy et al., 1999). Taking this 4-cycle delay into consideration, we reexamined the relationship between PSM length and somite size (comparing the size of the Nth somite with the PSM size at the N-4 ss, Fig. 1E). Strikingly, we found that somite size indeed scales with PSM size when this 4-cycle delay is considered (Fig. 1F). No clear relationship between somite and PSM length is apparent without the delay (Fig. 1F). This relationship between PSM length and somite size was still observed with a 3 or 5 cycle delay, suggesting that minor fluctuations in the delay or measurement error would not affect the conclusion (Fig. 1F). The delay between somite specification and formation is reflected in different peak positions in time course measurements of PSM and somite size (Fig. 1G). Consideration of this delay may be necessary to assess scaling in previous data (Gomez et al., 2008; Schroter et al., 2008).

**Fig. 1.**
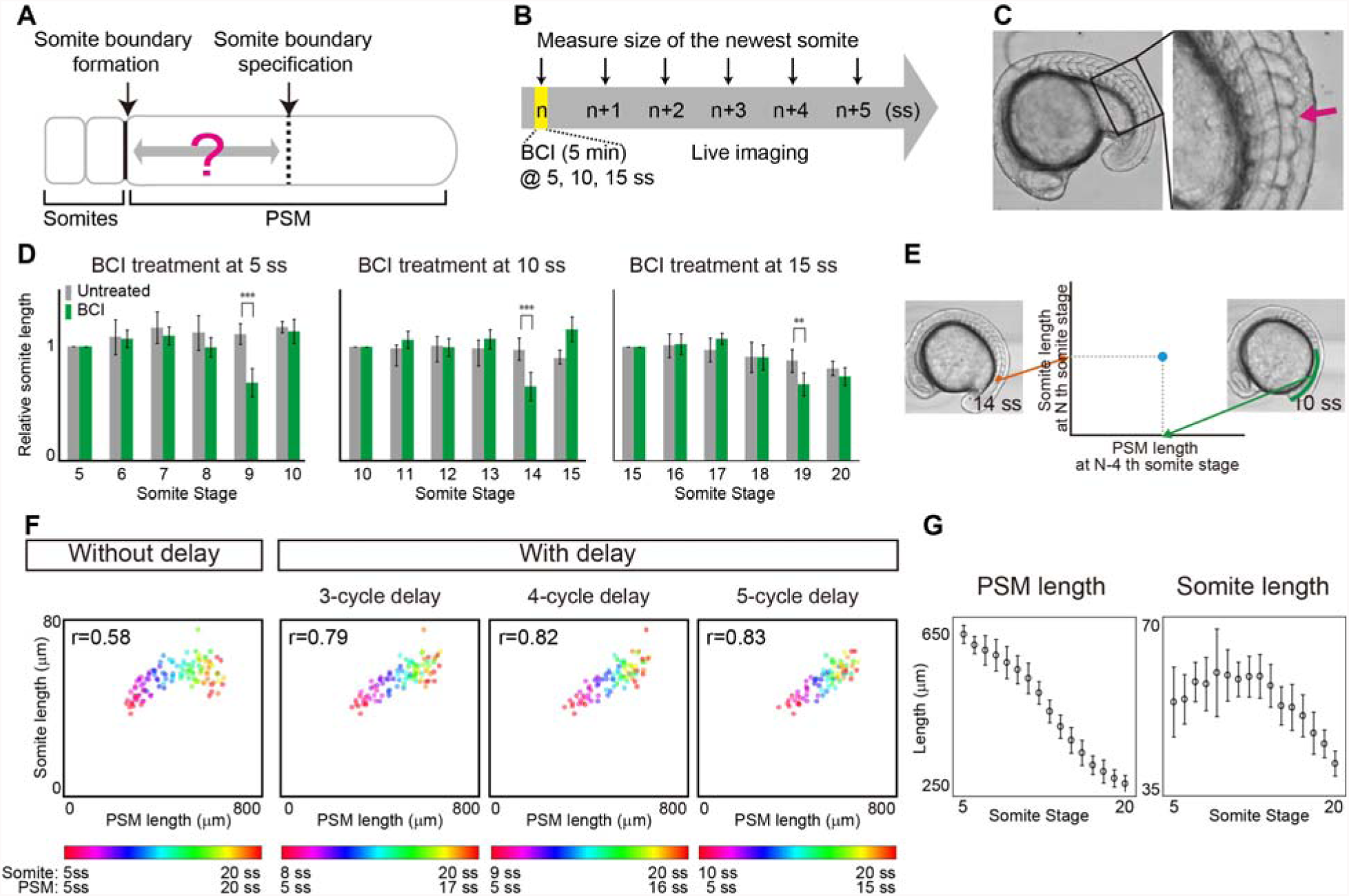
Somite scaling over time with time delay. (**A**) Schematic illustration of time delay between somite boundary specification and somite boundary formation. (**B**) Schematic illustration of BCI experiment. The embryos were treated with BCI for 5 min and then subjected to live imaging in egg water without BCI. The BCI treatment was done at 3 different somite stages (5, 10, 15 ss), in case the delay time varies over time. (**C**) BCI treated embryos form smaller somites (magenta arrow). (**D**) Relative AP length of somites, normalized by the somite length of control embryos at somite stage of BCI treatment. At each somite stage, the smaller somite was formed 4 cycles after BCI treatment. Error bars denote s.d.. **P < 0.01 and ***P < 0.001. (n=5 for each condition) (**E**) Comparison of PSM length and somite length was made using PSM length at N-4 ss (e.g. 10 ss) and somite length at N ss (e.g. 14 ss), using live imaging data. (**F**) Somite size vs PSM size at different somite stages with and without time delay (3, 4, 5 cycles). (**G**) Size dynamics of PSM and somites. Note the peaks appear at different somite stages. Error bars denote s.d.

### Somite length at specification scales with PSM length among individuals with different body sizes

Given that somite size at specification scales with PSM length throughout developmental time, we then wondered whether somite length scales with PSM length between zebrafish embryos of varying sizes. Inspired by classic work in *Xenopus* (Cooke, 1975) on somite scaling to body size in surgically size reduced embryos, we sought to apply this technique to zebrafish. We first attempted to cut zebrafish embryos at the blastula stage longitudinally (along the animal-vegetal axis) as was done in *Xenopus*. However, the resulting embryos had varying degrees of dorsalization or ventralization presumably due to dorsal determinants being portioned in unpredictable ways and were difficult to study quantitatively. We thus sought a method to reduce embryo size without perturbing D-V patterning. By using separate latitudinal cuts to remove cells near the animal pole and yolk near the vegetal pole at the blastula stage (Fig. 2A left panel), we found that the resulting size-reduced embryos quickly recovered and a large percentage of them developed normally (Fig. 2A). Total body size and organ size, including somites, of these size-reduced embryos were found to be smaller (Fig. 2B and C). Consistent with previous work in *Xenopus* (Cooke, 1975), the chopped embryos had the same number of somites, each of which was smaller in size (33 in both control and chopped embryos at 1 day post-fertilization, n=5 for each. Somite number was counted using still images of the live embryos). Combining this size reduction technique and live imaging, we measured somite and PSM length, and found somite length scales with PSM length between embryos of varying sizes when the same 4-cycle delay is considered (Fig. 2D, see also Fig. S2). The scaling was observed throughout our timecourses (from 5 ss to 20 ss, Fig. S3). Taken together, we conclude that somite length always scales with PSM length as long as the time delay between specification and morphological boundary formation is considered.

**Fig. 2.**
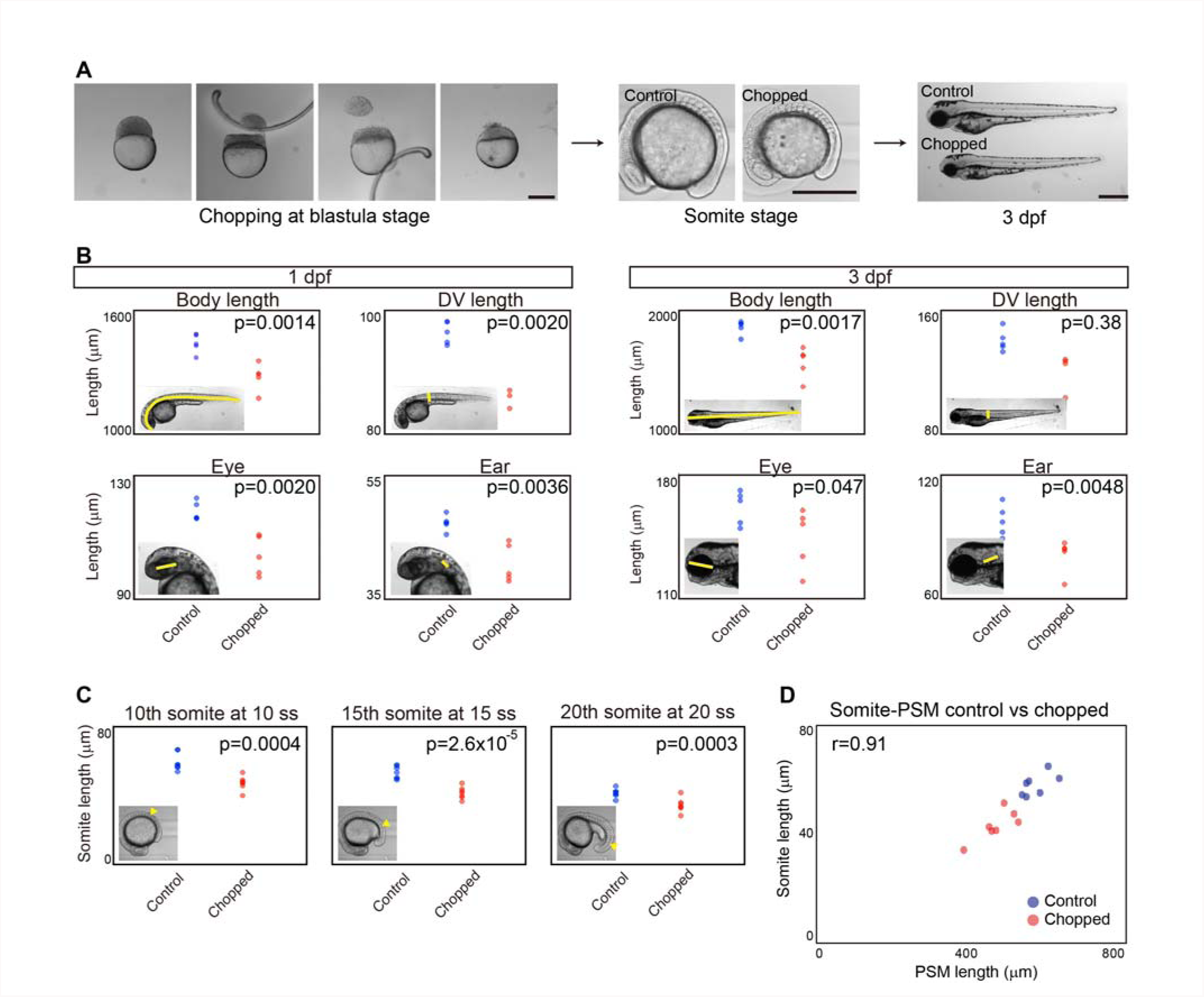
Somite scaling between individuals of different sizes. (**A**) Size reduction technique. Scale bar, 500 μm. (**B**) Body and organ sizes comparison between control and chopped embryos. (C) Somite size comparison between control and chopped embryos. (**D**) Somite size vs PSM size between control and chopped embryos.

### Clock period does not scale with PSM length

Given our finding that somite length scales with PSM length both over time and among individuals with different sizes, we next asked what mechanism might link PSM size to somite size. For this purpose, we searched for a component of the known somite patterning system that scales with PSM length, both across developmental stages and among individuals. In the classic clock and wavefront model, somite length is the product of clock period and wavefront regression speed. We first measured the period of the segmentation clock both in control and chopped embryos over time, since it is known that a change in the period of clock gene expression causes a change in somite length (Herrgen et al., 2010; Kim et al., 2011; Schroter and Oates, 2010). We measured the clock period as the time between the formation of successive somite boundaries, and found that this period is similar and does not scale with PSM length between control and chopped embryos (Fig. 3A, Fig. S4) or between those at different developmental stages (Fig. 3B) (Schroter et al., 2008), suggesting that scaling is not achieved by regulation of clock period.

**Fig. 3.**
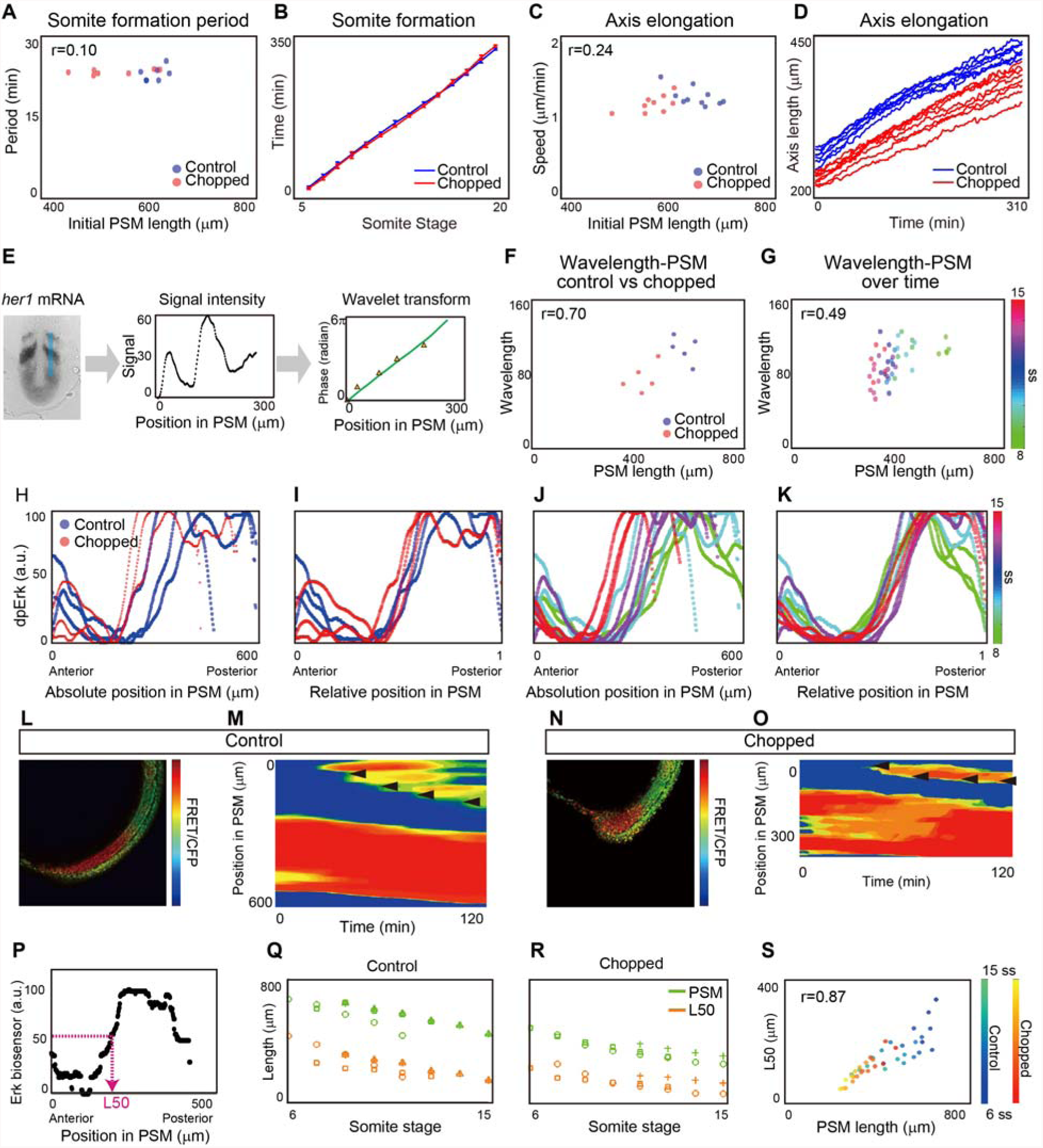
Determining which components of the somite formation system scale. (**A**) Somite formation frequency vs. initial PSM size. No significant difference was found between control and chopped embryos at the 5% significance level, and the confidence interval on the difference of means (−1.78 - 0.66) includes the hypothesized value of 0. (**B**) Somite formation time of control and chopped embryos. The slope corresponds to somite formation period. Note the slopes do not change over time. Error bars denote s.d. (**C**) Axis elongation speed vs. PSM size. No significant difference was found between control and chopped embryos at the 5% significance level, and the confidence interval on the difference of means (−0.10 - 0.15) includes the hypothesized value of 0. (**D**). Axis length vs. time. The slope represents the speed of axis elongation. (**E**) Quantification of *her1* wavelength along the blue line in the first panel. Green line in the third panel shows the phase gradient obtained by wavelet transform. Orange triangles show manually measured wavelength. (**F**) Wavelength vs. PSM among individuals. (**G**) Wavelength vs. PSM size over time. (**H** to **K**) Quantification of Fgf activity based on dpErk immunostaining. (**H** and **I**) dpErk scaling between control and chopped embryos. (**J** and **K**) dpErk scaling across developmental stages. Both are shown by absolute position (**H** and **J**), and relative position (**I** and **K**). (**L** to **S**) Quantification of Fgf activity based on Erk biosensor mRNA-injected embryos. The manipulated embryos (**L** and **N**) were used to generate kymographs of Erk activity (**M** and **O**). Black arrowheads represent newly formed somites. LUT, high (red) to low (blue) reporter intensity. (**P**) Definition of L50. (**Q** and **R**) Change in PSM size and L50 position overtime, in control embryos (**Q**) (n=4) and chopped embryos (**R**) (n=3). Different marks correspond to different embryos. (**S**) L50 vs PSM length over time both in control and chopped embryos.

### Axis elongation speed does not scale with PSM length

We next quantified the axis elongation speed, since slower axis elongation is known to lead to shorter somite length (Goudevenou et al., 2011; Rauch et al., 1997). One explanation for this comes from the clock and wavefront model, in which the wavefront speed (and hence somite size) has often been directly linked to axis elongation speed (Cooke and Zeeman, 1976; Dubrulle and Pourquie, 2004; Hubaud and Pourquie, 2014; Saga, 2012). This possibility is also consistent with the idea that a gradient of Fgf is established by mRNA decay coupled with axis elongation, and that this drives wavefront progression (Dubrulle and Pourquie, 2004). Therefore, we expected somites to be smaller in chopped embryos due to a decrease in the axis elongation speed (e.g. cells are incorporated into the PSM at the tailbud at a slower rate). We measured the change in axis length, defined by a distance between the posterior boundary of 4^th^ somite and the tail tip, over time (Bajard et al., 2014). Contrary to our expectation, we found that axis elongation speed did not differ between control and chopped embryos, at least for 5ss – 15ss (Fig. 3C, Fig. S4). This seemingly confusing result can be explained if the major mechanism of axis elongation at these stages is, for example, convergence and extension, whose rate should not be size dependent (Steventon et al., 2016). Notably, the axis elongation speed was nearly constant over our experimental time window (Fig. 3D), although PSM size decreased drastically. Since axis elongation speed neither changes over time as somites decrease in size nor between embryos of varying sizes, altered axis elongation speed cannot explain scaling of somite patterning.

### Wavelength of *her1* traveling waves does not scale with PSM length

We then asked if the wavelength of the traveling wave pattern of a segmentation clock gene (e.g. *her1*) could explain scaling of somite formation. Canonical segmentation clock genes exhibit traveling waves; a stripe pattern that sweeps through the PSM from posterior to anterior due to a phase delay toward the anterior direction. While these traveling waves have not been experimentally shown to cause somite size alterations, a correlation between wavelength (spatial interval of the stripes) and somite length has been observed (Jorg et al., 2016; Lauschke et al., 2013). To determine whether *her1* traveling waves are involved in scaling, we generated and quantified phase maps from *her1 in situ* hybridization samples (Fig. 3E). We extracted the phase information from signal intensities using a wavelet transform, then converted the approximately linear phase gradient into an effective wavelength, defined as the distance between peaks of *her1* intensity (Fig. 3E). We measured the phase gradient from an area of PSM including B-4 (the presumptive position corresponding to a morphological boundary four cycles later, blue line in Fig. 3E, left panel). We also measured the phase gradient manually, by identifying peaks and troughs in the intensity profile (orange triangles in Fig. 3E, right panel). This manual measurement was found to correspond well with phases obtained from the wavelet transform (green line in Fig. 3E, left panel). We found that unlike somite size, wavelength does not always scale with PSM size: although the wavelength scales with PSM size following embryonic size reduction, it does not scale during embryonic development (Fig. 3 F and G) (Holley et al., 2000). This is consistent with recent work demonstrating that the number of *her1* waves changes over time, confirming that the phase gradient does not scale with PSM size (Soroldoni et al., 2014). Since somite size scales with PSM size over developmental stages as well as among individuals of different size, this result indicates that it is unlikely that the somite scaling is achieved through regulation of the wavelength of *her1*. The conclusion is supported by a previous study which showed that repeated induction of *deltaC* expression in a *deltaC* mutant background can successfully rescue somite boundary formation, although the induced *deltaC* expression did not show the traveling wave pattern (Soza-Ried et al., 2014).

### The Fgf activity gradient scales with PSM length

Our final candidate feature that could relate somite size to PSM size was the Fgf gradient (Akiyama et al., 2014; Dubrulle et al., 2001; Sawada et al., 2001). To measure Fgf signaling we used whole mount immunohistochemistry against doubly phosphorylated Erk (dpErk), a downstream readout of Fgf activity, and extracted the signal intensity. We found that the gradient range varies considerably between embryos on an absolute length scale, but is consistent when plotted as a function of relative PSM length, both for control and chopped embryos (Fig. 3H and I, Fig. S5 and S6) and for embryos from different developmental stages (Fig. 3J and K, Fig. S5 and S6). We further tested if Fgf activity scales with PSM size in embryos carrying a FRET-based Erk biosensor (Fig. 3 L-S). We calculated the PSM location where the relative intensity of FRET signal crosses 50% of the maximal intensity (L50) (Fig. 3P). Recent work using the Erk live reporter showed that this L50 is a good approximation of the future somite boundary (Sari et al., 2018). Time course analysis of L50 in both control and chopped embryos confirmed that the Fgf activity gradient scales with PSM size (Fig. 3Q-S). L50 analysis was further performed when Fgf activity was measured by dpErk and by *sprouty4* mRNA, a downstream gene of Fgf signaling, and also confirmed Fgf activity scaling (Fig. S7). Since Wnt signaling is also known to form a gradient in the PSM, we examined whether Wnt signaling scales with PSM length. We performed L50 analysis on expression patterns of *sp5l* mRNA, a downstream gene of Wnt signaling (Thorpe et al., 2005), and found Wnt activity also scales with PSM length (Fig. S8).

### A clock and scaled gradient model can explain somite scaling

Given our observation of a dynamically scaling gradient, we turned to modeling to see whether this feature is capable of explaining scaling of somite patterning. In the original clock and wavefront model, the timing of somite boundary specification is controlled by a clock and the positioning by the level of a signal that encodes a posteriorly moving wavefront. How the position of the wavefront is determined at each time point is unspecified in the original model. Importantly, our observations reveal that the activity of signaling molecules linked with wavefront activity forms a dynamic gradient that scales with PSM size. We term this updated model the “clock and scaled gradient” model. In this model, scaling of the gradient to PSM size generates a posteriorly moving wavefront, when it is combined with axis elongation (which increases PSM size) and somite formation (which decreases PSM size) (Fig. 4 A and B). We constructed a simple mathematical model to formalize these interactions (Supplementary Materials and Methods) and found that this model can successfully reproduce our biological results on somite size scaling (Fig. 4C-F and Supplementary Movie). Similar somite formation dynamics can be observed regardless of the precise shape of the gradient (Fig. 4C and D; steep sigmoidal gradient, Fig. 4E and F; linear gradient). Interestingly, we also observed step-wise rather than continuous regression of the L50 in our model (Fig. 4G and H), consistent with recent results (Akiyama et al., 2014). Moreover using this model, we can also accurately predict the resulting changes in somite size following a wide range of additional previously-published perturbations (Fig. 4 I and J): one smaller somite following transient Fgf activation (Akiyama et al., 2014) (Fig. 4I); multiple smaller somites followed by one larger somite after Fgf bead transplantation (Dubrulle et al., 2001; Sawada et al., 2001) (Fig. 4I); larger somites with a slower clock (Herrgen et al., 2010; Kim et al., 2011; Schroter and Oates, 2010) (Fig. 4I); smaller somites with slower axis elongation (Goudevenou et al., 2011; Rauch et al., 1997) (Fig. 4I); and scaling of somite and PSM size *in vitro* under culture conditions that do not permit axis elongation (Lauschke et al., 2013) (Fig. 4J). We found that in all cases, the model’s predictions were in agreement with experimental results.

**Fig. 4.**
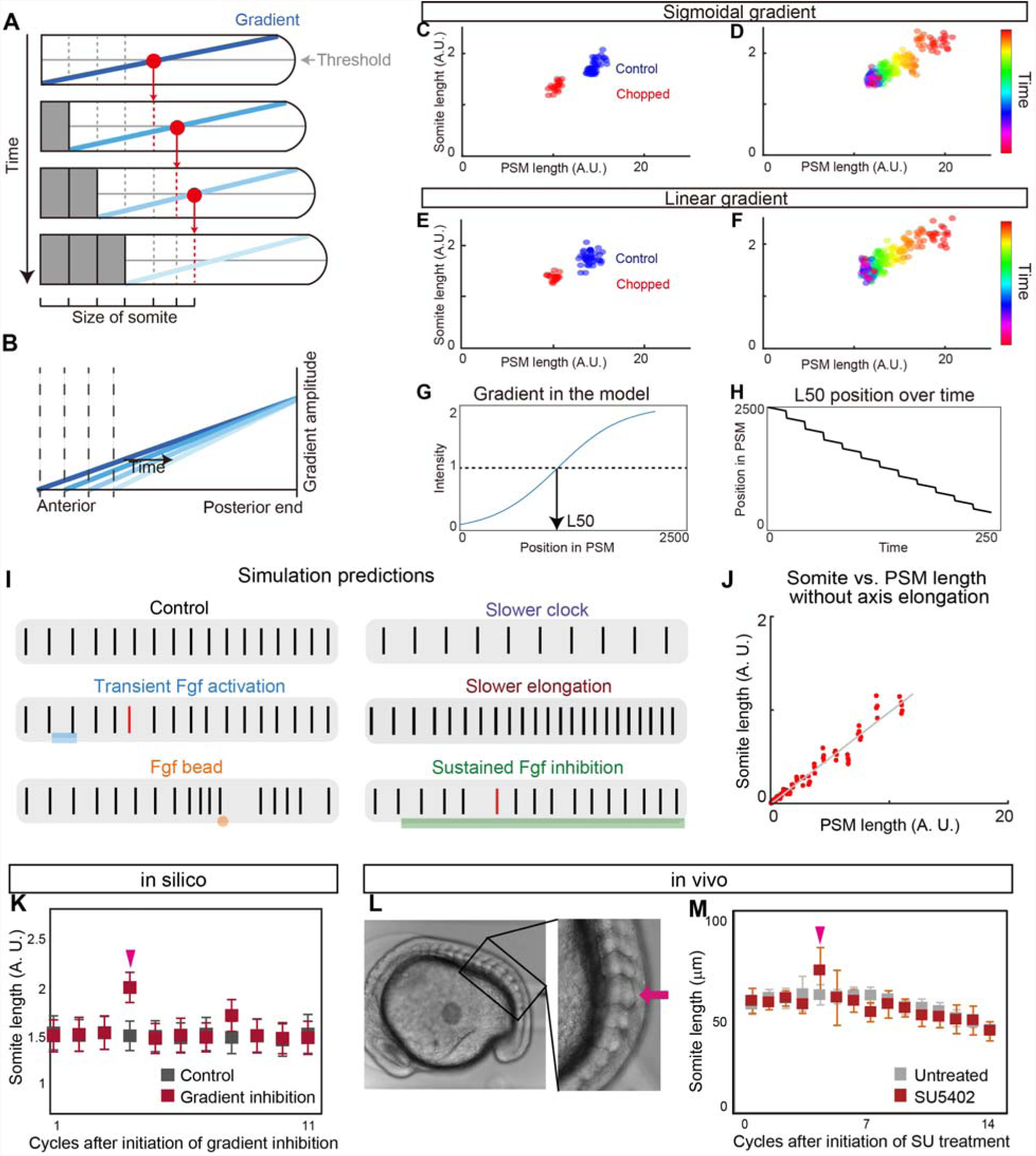
Clock and scaled gradient model. (**A**) Schematic illustration of the clock and scaled gradient model. (**B**) Superimposition of the gradients from each time point in (**A**). (**C** and **D**) Simulation results using a sigmoidal gradient. (**E** and **F**) Simulation results using a linear gradient. (**C** and **E**) Simulation results of control and chopped embryos. (**D** and **F**) Simulation results of a single embryo over time. (**G** and **H**) Stepwise regression of the gradient in clock and scaled gradient model. (**G**) L50 in the model was determined similarly to Fig. 3P. (**H**) Clock and scaled gradient model predicts stepwise regression of L50 position. (**I**) Simulation results for perturbation experiments for local or global inhibition/ activation of Fgf, slower clock and slower axis elongation. (**J**) Somite size versus PSM length shows perfect scaling *in silico* when axial elongation speed is zero, mimicking the results from the in vitro mPSM system (Lauschke et al., 2013). (**K**) Simulation results of long-term suppression of a gradient in the clock and scaled gradient model. Error bars denote s.d. (**L** and **M**) Low concentration of SU5402 (16 μM) results in one or two larger somite(s) (n=7 for both SU5402 and untreated). Error bars denote s.d.

### The clock and scaled gradient model predicts one larger somite in long-term Fgf inhibition

A simple perturbation to test our model is long-term Fgf inhibition. This experiment was recently carried out using chick embryos and multiple larger somites were shown to form during long-term Fgf inhibition (Cotterell et al., 2015). This result was contradictory to what the clock and wavefront model would predict, but consistent with a novel Turing framework for somitogenesis (Cotterell et al., 2015). We simulated the same perturbation using our clock and scaled gradient model and found that it predicts the same result as the clock and wavefront model: only one larger somite (Fig. 4K). To test if the long-term Fgf inhibition has the same effect in zebrafish embryos, we treated zebrafish embryo with the Fgf inhibitor, SU5402 (Sawada et al., 2001), at a low concentration (16 μM) in which embryos grew until late stages. Unlike in chick (Cotterell et al., 2015), we observed one larger somite but not multiple larger somites following long-term SU5402 treatment (Fig. 4L and M, Fig. S9, for individual data, see Fig. S13), consistent with our model. Moreover, we observed the same tendency under constant darkness, confirming the result we obtained was not due to the light instability of SU5402 (10 out of 11) (Fig. S10). These differences in results could potentially be explained by how acutely the drug can be administered: in zebrafish, embryos can be soaked in a vast excess of drug causing a rapid step up in drug levels followed by a plateau in vivo, whereas in chick the drug levels may rise more slowly. Simulations showed that increasing Fgf inhibition over a few hours can cause multiple large somites in our model (Fig. S11).

### Newly formed somites play a critical role in gradient scaling

One potential mechanism of gradient scaling is that newly formed somites modulate the gradient, for example, by secreting a negative regulator. To examine whether the newly formed somite can modulate the gradient, we transplanted a newly formed somite into the posterior PSM, and compared it to a control experiment in which PSM cells were transplanted to the same axial level (Fig. 5A). From our model, we predicted that the ectopically transplanted somite would locally inhibit Fgf signaling. One to two cycles (0.5-1 hour) after transplantation, the embryos were fixed and stained for dpErk. We found that in the PSM surrounding the transplanted somite, the dpErk level was significantly decreased (Fig. 5B), whereas the dpErk level in the PSM surrounding transplanted PSM cells was largely unaffected (Fig. 5C). To quantify Erk activity, we normalized the dpErk signal near the transplant with that of the non-transplanted side of the same embryo at the same axial level (Fig. 5A). We found the dpErk levels around the transplanted somite to be significantly lower than the control (Fig. 5D). These data support our hypothesis that mature somites rapidly and potently modulate the Fgf activity gradient to effect gradient scaling.

**Fig. 5.**
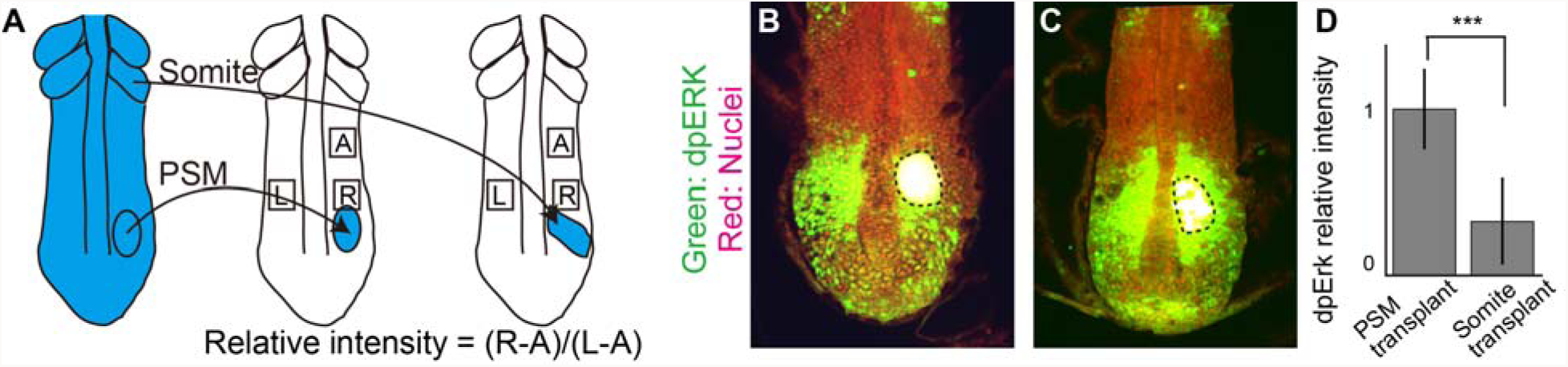
New somites inhibit Fgf activity. (**A**) Schematic illustration of somite transplantation. (**B** and **C**) dpErk immunostaining. Dashed line shows transplanted tissue. (**D**) Comparison of relative intensities between PSM transplanted samples (n=9) and somite transplanted samples (n=9). Error bars denote s.d. ***P < 0.001.

### A unique prediction from the clock and scaled gradient model: an “echo effect” on somite size

We next sought a novel experimental test for which our model makes a unique prediction. Key aspects of the clock and scaled gradient model are the 4-cycle delay between somite specification and formation, and the feedback between newly formed somites and gradient length. We thus reasoned that if we experimentally created one larger somite, it would shorten the PSM and rescale the gradient in a jump, which would then result in another larger somite four cycles later, and this process would repeat creating “echoes” of larger somites with a ~4-cycle periodicity (Fig. 6A). Simulations of our model supported this idea (Fig. 6B and C). To test this prediction we transiently treated embryos with the Fgf inhibitor, SU5402, which is known to induce a larger somite (Dubrulle et al., 2001; Sawada et al., 2001), followed by extensive washes for two hours, then performed live imaging to measure the length of the newly formed somites (Fig. 6D). Strikingly, we found that somite size became smaller and larger with a several-cycle period, which was uniquely predicted by the clock and scaled gradient model (Fig. 6E and F, for individual data, see Fig. S12). We noted that the periodicity was not always precisely 4 (Fig. 6F), possibly due to internal fluctuation of the delay time or experimental variation, such as variation in washout timing of SU5402. By analyzing individual embryos (Fig. S12), we confirmed that all the peaks of somite size in pulse SU5402 treated embryos are larger than those in control embryos (Fig. 6G). Our model also predicts the echo effect for long-term SU5402 treated embryos which we experimentally confirmed (Fig. S13), but we chose to focus on transient treatment because the embryos are healthier. The echo effect was also seen in embryos transiently treated with BCI (Fig. S14). These results confirm that the echo effect is a general phenomenon for somite formation. We note that a potentially related phenomenon has been seen following heat-shock in chick and zebrafish (Primmett et al., 1988; Roy et al., 1999) but through an unclear mechanism.

**Fig. 6.**
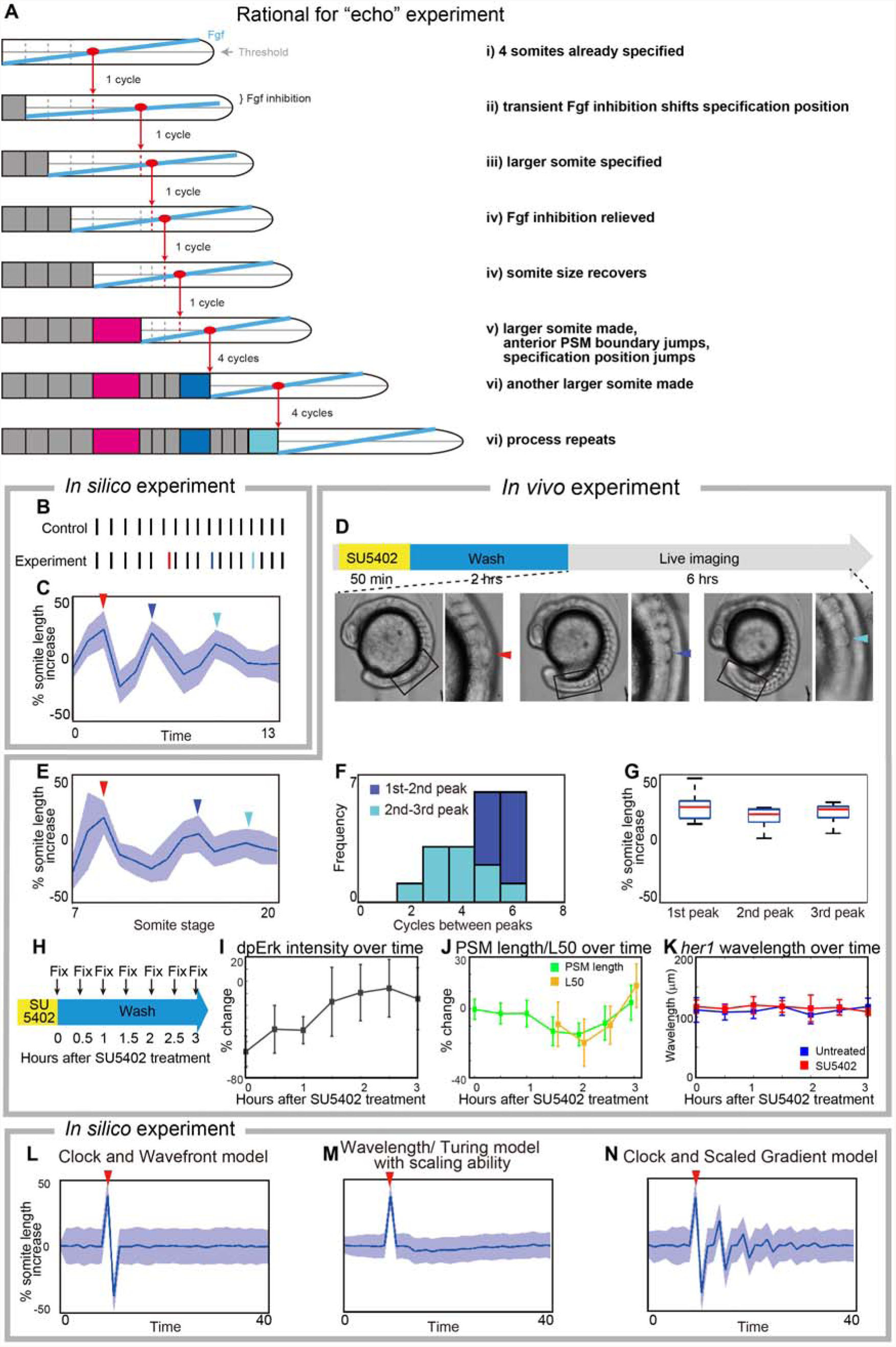
Experimental validation of the clock and scaled gradient model. (**A**) Schematic illustration of the outcome of the clock and scaled gradient model, following induction of one larger somite. The induced larger somite is colored in magenta, the larger somites as a result of system response are colored in blue and cyan. (**B** and **C**) Simulation results without (**B**) and with noise (**C**) for the somites size (red line in **B,** red arrow head in **C**). Blue line in **B** and blue arrowhead in **C** show the second, and cyan line in **B** and cyan arrowhead in **C** show the third large somite. (**D**) Schematic illustration of the *in vivo* experiment, and an embryo with larger somites at different time points. (**E**) Time course of percentage increase in somite length of SU5402 treated embryos, compared to those in control embryos (n=12). (**F**) Frequency distribution of somite cycles between the peaks. (**G**) Percentage increase in somite size in SU5402-treated embryos at the peaks detected in each embryo, compared to control embryos at the corresponding somite stage. In both **C** and **E**, blue lines and blue shades indicate the average somite size and the variance of one standard deviation, respectively. For **C**, **D** and **E**, red, blue and cyan arrowheads show the first, second and third larger somites. (**H-K**) Examination of Erk activity and *her1* wavelength after transient SU5402 treatment. Error bars denote s.d. (**H**) Schematic illustration of the experiment. After fixation, the samples were subjected to dpErk immunostaining and *her1* in situ hybridization. (**I**) Time course of percentage change in dpErk maximum intensity in SU5402-treated embryos, compared to those in control embryos. (**J**) Time course percentage change in PSM size and L50 position in SU5402-treated embryos, compared to those in control embryos. Note L50 analysis begins 1.5 hours after SU5402 treatment when dpErk intensity is recovered (see Fig. 6I) because it cannot be defined earlier. (**K**) Time course analysis of *her1* wavelength of untreated embryos and SU5402 treated embryos. We found no significant difference at significant level of 0.05 at any time point. (**L-N**) Simulation results for percentage increase of somite size over time, based on different models. After induction of one larger somite (arrowheads in magenta), the clock and wavefront model (when wavefront speed is associated with axis elongation only) predicts one smaller somite (**L**), wavelength/ Turing model (Cotterell et al., 2015) predicts smaller somites and the somite size eventually comes back to normal (**M**). Only the clock and scaled gradient model predicts the “echo effect” that somite size dynamics shows ups and downs repeatedly every 4 cycles (**N**).

We next evaluated the effect of transient SU5402 on both dpErk activity and *her1* wavelength (Fig. 6H-K, for individual dpERK data, see Fig. S15). To perform time-course analysis, we fixed the embryos every 30 min following SU5402 treatment (Fig. 6H). dpErk immunostaining confirmed quick recovery of Fgf activity after SU5402 treatment (Fig. 6I). Furthermore, as we assumed, the dpErk activity was found to scale with the induced smaller PSM (Fig. 6J). In contrast, we found no significant difference in *her1* wavelengths between control and SU5402 treated embryos (Fig. 6K), suggesting that gradient scaling, and not *her1* waves, are responsible for the echo effect.

This echo effect is only predicted if the “specification position” of new somites (rather than the somite itself) scales with PSM size, which is the core assumption of the clock and scaled gradient model (Fig. 6L-N). Without gradient scaling, the clock and wavefront model predicts a single smaller somite following the induced larger somite, but the size of the following somites immediately returns to normal (Fig. 6L), consistent with previous theoretical work (Baker et al., 2006). Interestingly, for a class of mechanisms that assumes that the “size” of a somite is what is determined, rather than the “position” of the next somitic furrow (e.g. somite size is determined by the wavelength of traveling waves, or the wavelength of a Turing-type pattern), then the predicted result is qualitatively different (Fig. 6M). In these models, somite size scales with the smaller PSM resulting from the induced larger somite, and then somite size gradually goes back to the normal size without rebounding dynamics. Together, the clock and scaled gradient model is uniquely supported by our experimental tests.

### Traveling waves have a minor effect in the clock and scaled gradient model

Spatial differences in the phase of the coupled oscillators comprising the segmentation clock give the appearance of traveling waves of clock gene expression in the PSM from the posterior to the anterior (Ares et al., 2012; Ay et al., 2014; Giudicelli et al., 2007; Morelli et al., 2009; Uriu et al., 2009), but a mechanistic role for these waves is unclear. Consistent with previous theoretical work (Ares et al., 2012; Morelli et al., 2009), our simulations show that traveling waves arise in systems of coupled oscillators under a wide variety of conditions including spatial variation in intrinsic frequency, coupling strength, and coupling phase delay, as well as due to differences in boundary conditions (Fig. S16), and thus their existence may not be significant. Thus far we have assumed synchronous oscillations throughout the PSM in our model for simplicity, as was done in the original clock and wavefront model (Cooke and Zeeman, 1976). To see if traveling waves affect the clock and scaled gradient model, we assumed a simple linear phase gradient along the AP axis (for details, see supplementary materials and methods) and repeated the simulations. As shown in Fig. 7A, this results in only a minor modification to somite sizes as compared to a model without a phase gradient. Interestingly, we noticed that the somite formation period (defined as the time between successive boundaries being specified) was smaller when including a phase gradient (Fig. 7B). This is consistent with the observation of the segmentation period in zebrafish being slightly faster than the intrinsic clock period - a phenomenon likened to the Doppler Effect (Soroldoni et al., 2014), in which an observer moving towards a source of traveling waves measures a higher frequency than the intrinsic frequency of the oscillators. We interpret this effect as caused by the wavefront moving towards the tailbud during development due to gradient scaling as the PSM shrinks rather than a change in arrival time of traveling waves to the anterior boundary (Fig. 7C). These results show that 1) phase-gradients have only minor effects on the clock and scaled gradient model and that 2) a model not based on traveling waves can also explain the Doppler effect.

**Fig. 7.**
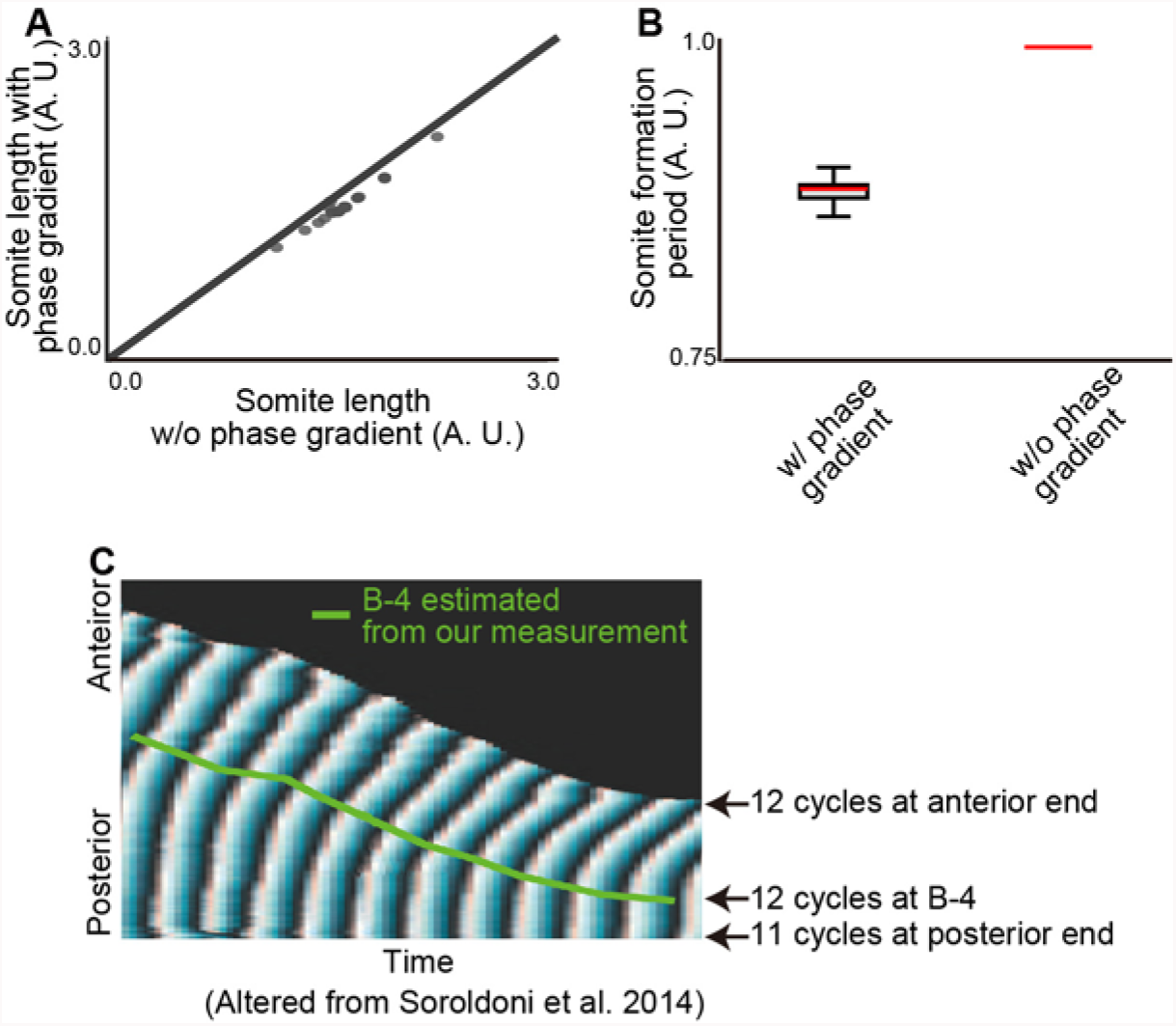
Traveling waves have modest effects on the clock and scaled gradient model. (**A**) Somite sizes are only slightly changed (~9%) by the presence of a phase gradient. (**B**) The phase gradient decreased the segmentation period (~11%). Error bars denote s.d. (**C**) B-4 positions estimated by our measurement are superimposed with the phase map generated previously (Soroldoni et al., 2014).

## Discussion

Here we have proposed a novel mechanism for somite size control: the clock and scaled gradient model. This model is based on the original clock and wavefront model but 1) the wavefront specifies new somite boundaries at a fixed *relative* position along the PSM due to gradient scaling, and 2) there is a delay between somite boundary specification and formation. Previously, multiple models of somitogenesis have been proposed, but were difficult to experimentally distinguish since they were all consistent with existing data from wild type embryos as well as existing experimental perturbations. Here we utilized a novel perturbation— changing system size—to discriminate between existing models, and showed that only the clock and scaled gradient model can explain existing data and our new experimental data. We found that in patterning of the somites, somite length scales with PSM length in vivo. Importantly, we demonstrate that the delay between somite boundary specification and formation is critical to examining the relationship between somite and PSM length. This is because the change in PSM length (and as a result, somite length) is dynamic, as a result of the changing rates of PSM production by axis elongation and consumption by somite formation (Fig. 1G). Consistently, when the PSM is grown in culture conditions that do not permit axis extension, there is a monotonic decrease in PSM size and somite-PSM scaling is observable without considering the delay (Lauschke et al., 2013). Considering the delay time between somite boundary specification and the appearance of a morphological somite will be essential in studying somite scaling in other situations, such as in other species, where complex dynamics of PSM length can be observed (Gomez et al., 2008; Schroter et al., 2008).

The clock and wavefront model is the classic model for somitogenesis (Cooke and Zeeman, 1976) and explains a number of previous experimental observations. In the original clock and wavefront model, what controls wavefront progression and how it is linked to axis elongation is unspecified. A simple way to specify wavefront progression is to just tie it to axis elongation (Cooke and Zeeman, 1976; Dubrulle and Pourquie, 2004; Hubaud and Pourquie, 2014; Saga, 2012). The consequence of this version of the clock and wavefront model is that somite size is equal to how far the tail moves in one clock cycle. While many of the existing perturbations can be explained by this version of the clock and wavefront model, a key feature that cannot be predicted is the phenomenon of scaling. For example, in the absence of axis elongation no somites should form, but this prediction is contradicted by in vitro cultured PSM which has no axis elongation yet forms a series of progressively smaller somites (Lauschke et al., 2013). Similarly, this simple clock and wavefront model does not predict the non-monotonic somite size variations following induction of a single large somite, as seen in Fig. 6, since perturbations to the anterior PSM should not affect wavefront position.

An alternative class of models for explaining somite formation is based on using the wavelength of traveling waves in determining somite size (Jorg et al., 2015; Jorg et al., 2016; Lauschke et al., 2013). However, previous studies in addition to our new results suggest that the wavelength of the traveling waves is not the primary mechanism to set somite sizes. First, the simple scenario (Lauschke et al., 2013) assumes that the phase gradient (inverse of wavelength) of the entire PSM scales with PSM length and that the scaled wavelength sets the somite size. However, previous results (Soroldoni et al., 2014) and our results show that phase gradient does not always scale with PSM length, which argues against this simple mechanism. Second, one could still imagine some modification of the simple wavelength model could explain the in vivo situation of somite scaling (e.g., the wavelength at B-4 locally scales with PSM length). However, this model is still hard to reconcile with the echo effect we observed after inducing one large somite (Fig. 6), because regardless of the details, this class of models assumes somite “size” (not somitic furrow position) is controlled by the wavelength. In Fig. 6M, we explicitly model the case where somite sizes scale with PSM size (including the 4-cycle delay) and find that it cannot explain the echo effect. In order to directly test if traveling waves are functional, one should experimentally modify the spatial pattern of the waves (for example, changing or eliminating the spatial phase gradient), without affecting the intrinsic period of the oscillators (Soza-Ried et al., 2014), and a mechanism for detecting a spatial gradient in clock gene expression level should be proposed. We suggest that traveling waves may be a byproduct of the need to synchronize oscillators locally (within the spatial scale of a somitic furrow), that while visually striking and mathematically interesting, have only a peripheral role in somite formation.

Another type of model is ‘Turing-like’, in which somites are formed via a combination of an oscillator and a periodic Turing instability (Cooke and Zeeman, 1976; Cotterell et al., 2015). There are several reasons why our data does not support Turing-like models. Firstly, a recent paper (Cotterell et al., 2015) showed how a Turing-like model of somitogenesis could, in principle, explain somite scaling, if one allowed the level of Fgf to effectively modulate the Turing-spacing of the somites. However, the change in somite size in response to PSM length is small, and is inconsistent with our in vivo measurement where somite length is almost proportional to PSM length (Fig. 1F and 2D). A second argument against a Turing-like model is that, unlike the clock and wavefront and clock and scaled gradient models, the ‘clock’ is not separable from the other components in the system. Therefore we don’t necessarily expect a slower clock to increase somite size, at least not in perfect proportion as has been observed in vivo (Herrgen et al., 2010; Kim et al., 2011; Schroter and Oates, 2010) since a change in clock period would be associated with other parameters. Thirdly, the assumption that Fgf modulates the Turing-spacing of somites is incompatible with the results of perturbing Fgf, specifically: 1) a Turing-like model predicts consistently larger somites following sustained Fgf inhibition, which we do not see (Fig. 4 L and M); 2) a Turing-like model predicts a symmetric effect of implanting a Fgf bead (i.e. smaller somites anterior and posterior to the bead) unlike what is seen in vivo which shows a definite anterior-posterior bias (Dubrulle et al., 2001; Sawada et al., 2001). Finally, it is difficult to reconcile a Turing-like model with the echo effect (Fig. 6). The reason is that, like the phase-gradient model, and unlike the clock and scaled gradient model, Turing-like models fundamentally control somite size, not somite boundary position. Therefore, for exactly the same reasons as argued for the phase-based models, even with perfect somite size scaling in wildtype embryos, we do not predict the non-monotonic segment size variation following transient Fgf inhibition.

The clock and scaled gradient model presented here is a fairly simple model. We used a simple model for three reasons: 1) so that the key assumptions of the model (clock + scaling gradient) are directly supported by experimental data; 2) so that the model is at the right level of detail to make comparisons to our data; and 3) so that the model gives us a qualitative and intuitive understanding of somite size control, which may be obscured in a more complex model (Gunawardena, 2014). However, the model’s simplicity does mean that it should not be viewed as a comprehensive, nor completely realistic, model of somitogenesis. Firstly, we have assumed that somite maturation, and its effects on gradient scaling, occur instantly, whereas in reality we expect this to be a more gradual effect. Mathematically, this might mean that the 4-cycle delay should be changed from a step function to a more slowly varying function. This modification may be particularly important to understand the formation of the first four somites, and to reduce the sensitivity of somite size to initial conditions and/or perturbations. A second shortcoming of our model is that we have chosen the somite boundary to be set by a simple threshold of the gradient - an assumption that has not been directly measured, and is likely a simplification. Thirdly, we have largely focused on dpErk as readout of wavefront activity and demonstrated dpErk scaling as a proof of concept. However, the wavefront could be set by a complex function of multiple inputs such as Fgf and Wnt along with downstream signal integration (Bajard et al., 2014; Stulberg et al., 2012; Wahl et al., 2007), without affecting the core conclusions of our model. As reported, dpErk shows a steep gradient (Akiyama et al., 2014), but in our model, similar somite formation dynamics can be observed regardless of the precise shape of the gradient; even a simple linear gradient can recapitulate the *in vivo* behavior rather closely (Fig. 4E and F). And finally, the molecular mechanism of gradient scaling remains to be determined. Numerous regulatory interactions have been shown in the posterior axis between Fgf, Wnt, Brachyury, Sprouty, and Retinoic Acid so these are all candidates (Diez del Corral et al., 2003; Olivera-Martinez and Storey, 2007).

One reason we chose to look at scaling of somites in size-reduced embryos is that we thought we might discover a mechanism for scaling that is *not* based on scaling of a molecular gradient (e.g. change in axis extension speed, growth rate, phase gradient, oscillation period). However, in the end we found that scaling of a molecular gradient is indeed what underlies somite scaling as has been observed in other examples of pattern scaling (Ben-Zvi and Barkai, 2010; Howard and ten Wolde, 2005; Umulis and Othmer, 2013). Future research on this issue could reveal what design benefits (e.g. robustness, evolvability) systems employing gradient scaling have compared to other potential mechanisms for scaling.

## Methods

### Fish care

Fish (AB) were kept at 27°C on a 14-hr-light/ 10-hr-dark cycle. Embryos were collected by natural crosses. All fish-related procedures were carried out with the approval of Institutional Animal Care and Use Committee (IACUC) at Harvard University.

### Size reduction technique

Chorions were enzymatically removed using pronase (Sigma Aldrich, 1mg/ml in egg water (Westerfield, 2000)) at ~512 cell stage. Eggs were treated with pronase until the chorions loses their tension and washed gently with egg water. Remaining chorions were removed manually using tweezers. The embryos were placed on a glass dish with 1/3 Ringer’s solution (Westerfield, 2000), with 2% methylcellulose (Sigma Aldrich) in 1/3 Ringer’s solution spread thinly on the bottom of the dish, to restrict movement embryos. We found using 1/3 Ringer’s solution is critical for embryos to recover from the damage of chopping. Then the blastoderm was chopped at the animal pole, and the yolk was wounded, resulting in oozing out of the yolk, using either hand-pulled glass pipette or looped stainless steel wire (30 μm in diameter) glued in the capillary glass. The chopped embryos were incubated in the 1/3 Ringer’s solution for 30 min, and then moved to fresh 1/3 Ringer’s solution for further incubation. The survival rate of the chopped embryos varies depending on condition of the embryos. Healthy embryos and good dissection would give ~60% success rate of developing normally until late stages (at least several days). The ratio of remaining cells and yolk affects how normal the embryos develop; usually cutting 50% position of blastula horizontally and wounding the vegetal part of yolk produces good results.

### BCI and SU5402 treatment

Embryos were treated with BCI (Dual Specificity Protein Phosphatase 1/6 Inhibitor, Calbiochem) as described (Akiyama et al., 2014). For SU5402 treatment, embryos were treated at a low concentration (Calbiochem, 16 μM) to minimize toxicity.

### Imaging

For live imaging, the embryos were mounted laterally using the dorsal mount (Megason, 2009) in egg water with 0.01% tricaine (Wentern Chemical, Inc.). Live imaging was performed using Zeiss Axio Observer Z1 and AxioCam MRm. For multiple image acquisition, we used a motorized stage, controlled by AxioVision 4.8. The temperature was maintained at 28.5 ± 0.5°C using a home-made incubator. The images were taken every 2 min, and the size of z slice varied depending on the size of embryos. The images of the *in situ* hybridization samples were also acquired using Zeiss Axio Observer Z1. The images of dpErk immunostaining samples were acquired using Leica TCS SP8. Finally, a Nikon Ti spinning disk confocal was used to acquire the images of transplanted samples.

### Image processing

Image processing was done using FIJI (Schindelin et al., 2012) and Matlab custom code. For time course measurement of axis elongation and somite size, we used the Gaussian-based stack focuser in FIJI. For axis elongation measurement, the length from 4^th^ somite to tail tip was measured, using FIJI’s LOI interpolator. For *in situ* hybridization samples and immunostaining samples, noise was first reduced using Gaussian blur (sigma = 7.0), and the signal was extracted along AP axis, using FIJI’s Plot profile function. To compare intensity profiles of BCI and SU treated embryos (Fig. S1 and S9), we averaged over multiple embryos. To calculate relative intensity, first, the minimum value was set to 0; and then the intensities at each position was scaled with a scaling factor of (average maximum intensity in drug treated embryo/average maximum intensity of untreated embryo).

### *In situ* hybridization and immunostaining

*In situ* hybridization (Nikaido et al., 1997) was performed as previously described. dpErk immunostaining was performed basically following the protocol described in Sawada et al.(Sawada et al., 2001), except that we used Alexa Fluor 488 goat anti-mouse IgG (ThermoFisher Scientific A-11001) as the 2^nd^ antibody. Nuclei were stained with propidium iodide (Life Technologies P1304MP).

### Somite/PSM transplantation

Transplantation was performed as described (Haines et al., 2004; Kawanishi et al., 2013), with minor modification. For making a cut on the skin, we used a mouth pipette filled with pancreatin, so the cut can be made both physically and enzymatically. Embryos for donor tissue were injected with Alexa Fluor 680 conjugated 10,000 MW Dextran, which can be detected directly after immunostaining.

### Live imaging of Erk activity dynamics

The FRET-based Erk biosensor termed Eevee-ERKnls is composed by an enhanced cyan-emitting mutant of GFP (ECFP), a WW domain (ligand domain), an EV linker, an Erk substrate (sensor domain), a yellow fluorescent protein for energy transfer (Ypet), and a nuclear localization signal (NLS) (Komatsu et al., 2011). When Erk phosphorylates the Erk substrate, the WW domain binds to the Erk substrate, leading to the induction of FRET from ECFP to Ypet. It has been confirmed that the Erk biosensor can monitor Fgf-dependent Erk activity in living zebrafish embryos (Sari et al., 2018). One cell stage of embryos were injected mRNA encoding a *FRET-based ERK biosensor* termed *Eevee-ERKnls* (Sari et al., 2018; Komatsu et al., 2011). The embryos at a certain stage were excited with a 440-nm laser, and fluorescence spectra were acquired by using a Lambda Scanning mode of a LSM710 confocal microscope (Zeiss). Using a Linear Unmixing mode, CFP and Ypet signals were separated from the original spectra data. FRET/CFP ratio images and kymographs were created with MetaMorph software (Molecular Device).

### Statistical test

Significance was calculated by one-tailed Student’s t tests, using Excel (Microsoft). Unequal variance comparison was performed for Fig. 1D, Fig. 2 B and C, and equal variance comparison was performed for Fig. 5D and Fig. 6K.

### Wavelet transform

We follow the approach of (Soroldoni et al., 2014) and use the wavelet transform to generate phase maps for *her1* along the embryo. Consider that the *her1* pattern is of the form:

*h(x) = h + A(x)sin(ϕ (x) +Φ* i.e. has a spatially varying amplitude, *A*(*x*) and a spatially varying phase, *ϕ*(x). By performing a wavelet transform we can convert the intensity profile *h*(*x*) into an effective phase profile *ϕ*(*x*), plotted in Fig. 3E. Note, we plot the phase for positions more anterior than the first clear peak since it is only in these ranges where there is a distinct spatial pattern above noise, and, in all cases, contains the position at which the next somite boundary is specified i.e. B-4. We also measured the phase gradient manually, by identifying peaks and troughs in the intensity profile (separated by π). This manual measurement (orange triangles in Fig. 3E) was found to well match the corresponding phases as obtained from the wavelet transform, giving us confidence in our implementation. For further details of the wavelet transform, we refer the reader to (Soroldoni et al., 2014).

## Acknowledgments

We thank O. Pourquié, A. Aulehla and A. Oates for critical discussion. Some images were taken at Nikon Imaging Center at Harvard Medical School. K.I. acknowledges a Postdoctoral Fellowship for Research Abroad (Japan Society for Promotion of Science).

## Author contributions

K.I. conceived the study and conducted experiments and data analysis. T.H. did modeling, simulations and wrote the program for wavelet analysis. Z.C. established size reduction technique with K.I. D.S., K.L. Y.B. and T.M. established Erk reporter fish and performed live imaging for this reporter. D.R. helped with image analysis. S.M. supervised the overall study. K.I., T.H. and S.M. wrote the paper with input from other authors.

